# Morphogen-driven human iPSCs differentiation in 3D *in vitro* models of gastrulation is precluded by physical confinement

**DOI:** 10.1101/2023.03.29.534685

**Authors:** Haneen S. Alsehli, Errin Roy, Thomas Williams, Alicja Kuziola, Yunzhe Guo, Jeremy Green, Eileen Gentleman, Davide Danovi

## Abstract

In early human development, gastrulation is tightly associated with lineage specification. The interplay between mechanical forces and biochemical signals during these processes is poorly understood. Here, we dissect the effects of biochemical cues and physical confinement on a 3D in vitro model of gastrulation that uses spheroids formed from human induced pluripotent stem cells (hiPSCs). First, we compare self-renewing versus differentiating media conditions in free-floating cultures, and observe the emergence of organised tri-germ layers. In these unconfined cultures, BMP4 exposure induces polarised expression of SOX17 in conjunction with spheroid elongation. We then physically confine spheroids using PEG-peptide hydrogels and observe dramatically reduced SOX17 expression, albeit rescued if gels that soften over time are used instead. Our study combines high-content imaging, synthetic hydrogels and hiPSCs-derived models of early development to define the drivers causing changes in shape and emergence of germ layers.

## Introduction

During the early stages of human development, the pluripotent stem cells of the epiblast undergo gastrulation, breaking symmetry to generate anterior-posterior axial elongation and shaping the body plan. This process is temporally associated with lineage specification; the pluripotent stem cells eventually give rise to all cell types in the mammalian body [1]. Mechanical interactions between these cells and the extra-embryonic environment are key regulators of morphogenesis and gastrulation events in mammals [2, 3]. However, the interplay between mechanical interactions that drive morphogenesis and lineage commitment during early development is difficult to study *in vivo* and poorly understood [3, 4].

Human induced pluripotent stem cells (hiPSCs) offer an unprecedented tool to study cell fate decisions [5], self-organisation, and early developmental events [5, 6]. The molecular signals regulating self-organisation during specification of the three germ layers have been investigated *in vitro* using a variety of models [7, 8]. Several studies have proposed that SOX17 is expressed within the inner mass cells (ICM) and in primitive endoderm (PE) concomitantly with lineage differentiation [9–11]. In other organoid systems [8, 12] the dynamic interactions between the intrinsic mechanical properties of the surrounding tissue [13], and the forces and soluble signals cells generate and receive affect cell behaviour [2, 7]. Nevertheless, further investigation is needed to understand how these processes are coordinated to influence cell fate decisions. Indeed, the requirement for morphogenesis (if any) to achieve proper differentiation has not been clearly established [14, 15].

Micropatterned systems in 2D have been used to investigate the signalling pathways underlying self-organisation and fate decisions in early human development. Warmflash *et al*. demonstrated that human PSCs cultured on defined patterns could radially organise and differentiate into cells of the three germ layers following BMP4 treatment, mirroring early development [1, 8, 16, 17]. Micropatterned colonies differentiated into ectoderm in the innermost region, surrounded by concentric rings of mesoderm and endoderm cells with an exterior trophectoderm ring [8, 16]. Building on these findings, it has been proposed that mechanical tension generated by cell-adhesion in 2D gastrulation models is a key regulator of cell fate specification [2, 18]. Here, different geometrical shapes (circle, square, and triangle) alter BMP4-mediated patterning in regions of high tension [2]. These approaches have advanced our knowledge of the role geometry plays in lineage commitment and signalling [8, 19]. Indeed, we further showed that these models are suitable to highlight relationships between genetic variations and germ layer differentiation [18] for selected hiPSCs lines (see [20]).

On the other hand, 3D cultures offer a more realistic model to investigate the role of physical forces in response to environmental cues in conjunction with morphogen-triggered signalling events. Various culture methods can establish embryo-like axial elongation and display key features including gene expression patterns mirroring normal development [1, 21, 22]. A recent study described that hESCs cultured in suspension under defined conditions treated with pulses of the Wnt agonist Chiron, drove symmetry breaking and elongated morphologies without the addition or formation of extra-embryonic tissues [1, 23]. The resulting ‘gastruloids’ displayed polarised expression of mesoderm (BRA), endoderm (SOX17) and neuroectoderm (SOX2) markers. These observations suggest that Wnt signalling is sufficient to initiate axial elongation and patterning of the three germ layers in 3D human iPSCs cultures. In contrast, under similar conditions, in the presence of BMP4 but without Wnt signalling, cells were reported to fail to aggregate, elongate and adopt noticeable patterning [1]. This supports the hypothesis that in addition to signalling, the physical/mechanical environment is crucial prior to and during lineage specification *in vitro* to achieve patterning and axial elongation [1].

Hydrogels can be used to manipulate intrinsic mechanical properties in 3D in vitro models of gastrulation [7, 24]. Such platforms can control matrix stiffness, degradability and cell adhesion and have been used to study the role intrinsic mechanical cues play in supporting intestinal organoids and controlling neural tube morphogenesis [25, 26]. Hydrogels provide well-defined, reproducible environments that control both physical and biochemical cues [27]. Indeed, hydrogel stiffness can be tuned by varying the polymer concentration, and biological cues controlled by incorporating integrin-binding (RGD) and matrix metalloproteinase-degradable peptide sequences [26]. Matrix stiffness is a key regulator for multiple cellular processes including proliferation, differentiation, migration and spreading. Importantly, previous work showed that intestinal organoid formation is favoured in hydrolytically degradable [28] or viscoelastic [29] (rather than purely elastic) matrices. In hiPSCs, adhesion signalling directs morphogenic events such as lumen formation and apicobasal polarity in 3D hiPSCs [30].

In this study, we investigated how biochemical cues and physical confinement separately influence morphogenesis and differentiation in a 3D hiPSCs model of gastrulation. Observing and quantifying how spheroid-forming cells behave within defined medium conditions upon BMP4 treatment, we specifically question whether morphogenesis is required to direct lineage differentiation and patterning. [31, 32] We hypothesised that BMP4 provides sufficient signals to trigger symmetry breaking, elongation and differentiation of hiPSCs spheroids in suspension culture. We also hypothesised that synthetic hydrogels [28, 33] inhibit elongation, thus enabling us to interrogate whether physical confinement affects not just morphogenesis, but also cell fate specification. Taking advantage of previous 2D micropattern systems [8, 11], we adapted a similar set up in our 3D model. When hiPSC spheroids were encapsulated within hydrogels, elongation was impeded and SOX17 expression dramatically reduced. Our approach enables us to investigate the inter-correlation of changes in shape with patterning of germ layers in hiPSCs-based models of early development.

## Results

### BMP4 signalling induces axial elongation in 3D gastrulation-like models

To quantify dynamic changes in the morphology of hiPSCs spheroids in suspension, we first set up a robust high-content imaging-based platform to monitor morphology using frame-to-frame variation analysis to obtain efficient segmentation of simple phase-contrast images and precise morphology quantification [34]. We used defined medium conditions that provided consistent shape variation in spheroid morphology. E8 medium promotes self-renewing conditions, while KSR BMP4 triggers differentiation [31, 32]. We monitored hiPSCs aggregates forming spheroids when cultured in these conditions and controls (Figure 1). Consistent with our previous results, we observed distinct phenotypic variations in each medium condition (Figure 2A-D). Cells cultured in E8 medium formed round spheres (movie S1), whereas in KSR BMP4, spheroids broke symmetry and exhibited axial elongation (movie S2). The other conditions produced intermediate phenotypes: in E8 BMP4, spheres were smaller in size, and few tended to form dynamic small protrusions (movie S3), while culturing in KSR alone produced a budded morphology (movie S4). These results demonstrate that the addition of BMP4 to KSR medium enabled consistent spheroids symmetry breakage by axial elongation.

**Figure 1.**
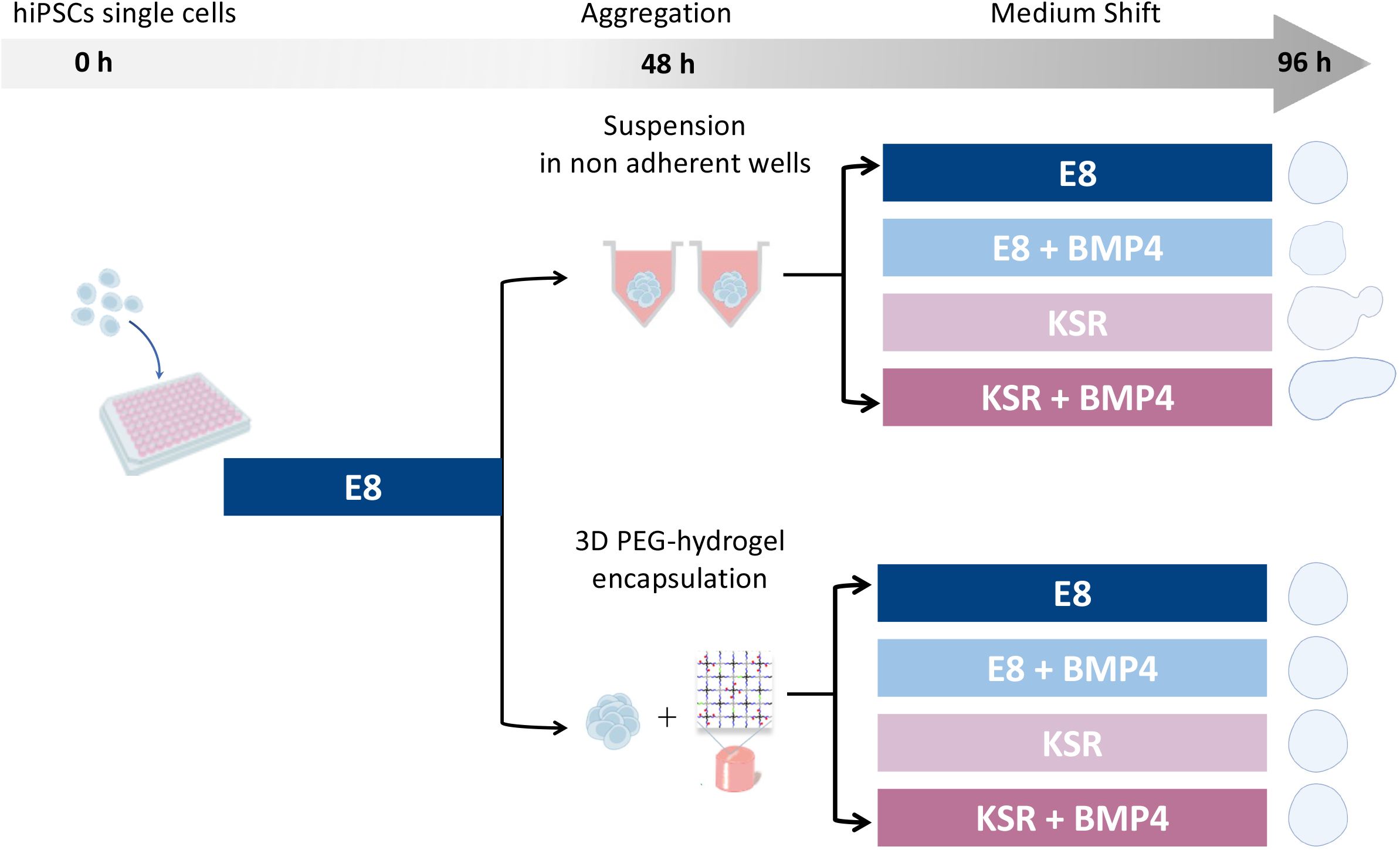
Schematic of the experimental workflow. hiPSCs are seeded in low attachment wells plate cultured in E8 medium for 2 days to form spheroids in suspension (top) or hiPSCs spheroids were encapsulated in PEG-4VS hydrogel (bottom). Medium is changed at day 2 either cultured in E8 medium to maintain pluripotency or KSR BMP4 to form gastruloids-like structure. E8 BMP4 medium and KSR alone are control conditions, spheroids are imaged every hour from day 2-6.

**Figure 2.**
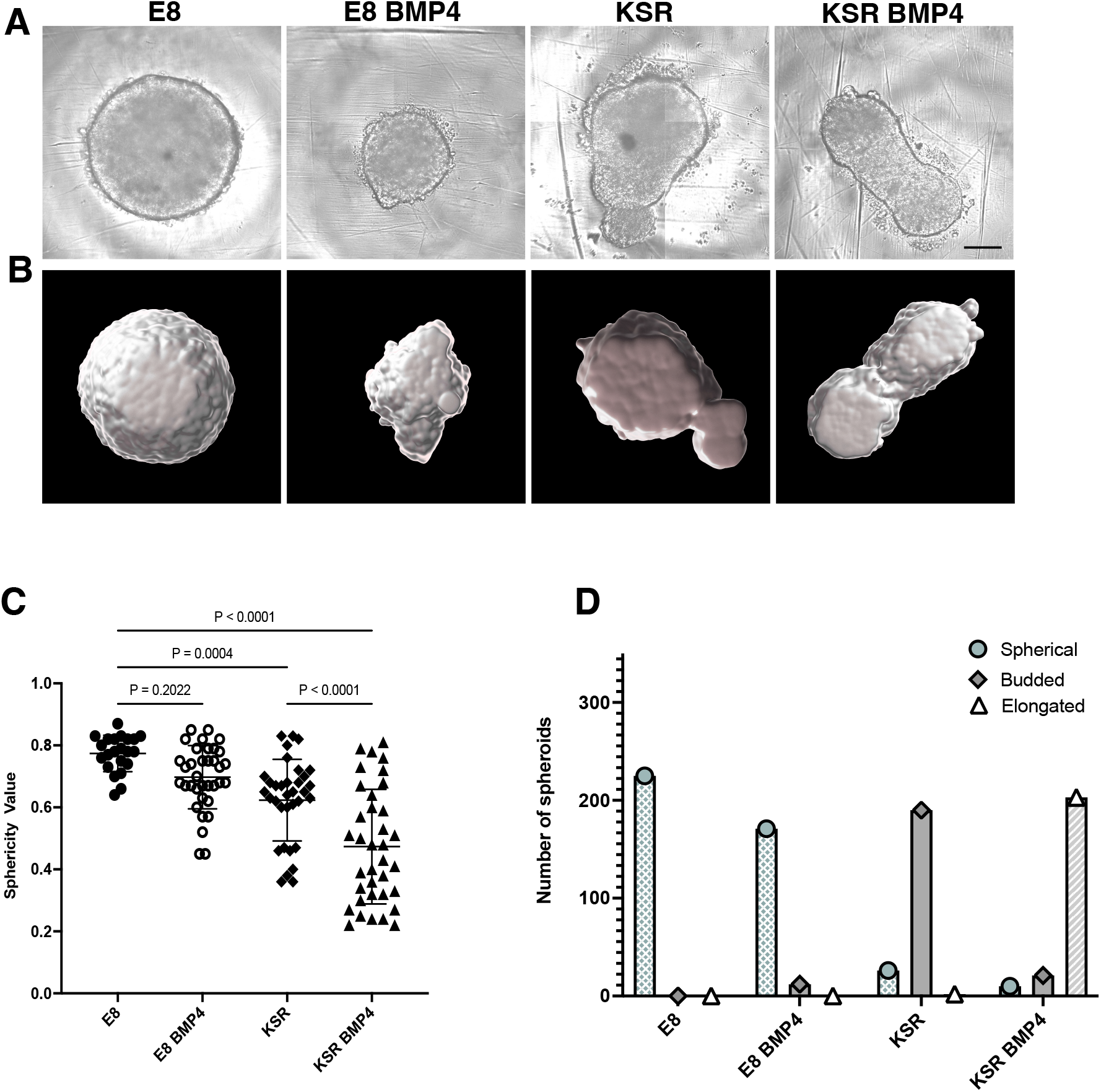
hiPSCs forming spheroids with a distinct morphological change in different medium conditions. **A)** The changes in spheroid shape, E8 medium maintained spherical shape, KSR BMP4 induced axial formation, in controls conditions E8 BMP4 small spheroid tended to form a budding, while KSR formed budded structure (Scale bar 200 um). **B)** Confocal representative images of 3D culture and spheroids were masked using Imaris software to quantify deformation in shape. **C)** Sphericity values were obtained from the segmented images, in E8 0.8 value indicates a round shape, in KSR BMP4 value varies between 0.2 – 0.7 which demonstrates an elongated shape, E8 BMP4 exhibits rounded values while KSR is more rounded compared to KSR BMP4 (n=22, 36 spheroids in E8, and rest of conditions respectively, and SD).

### Morphogenesis drives SOX17 patterning upon BMP4 treatment

To confirm differentiation concomitant with axial elongation, we first stained for pluripotency marker OCT4. We found that spheroids cultured in E8 medium expressed OCT4, while it was expressed at only low levels in cells cultured in the other three conditions (Figure 3A-B) (P value all <0.0001). Notably, the KSR BMP4 medium, despite its ability to drive the most substantial shape change, left some residual OCT4 expression, which co-expressed with SOX17.

**Figure 3.**
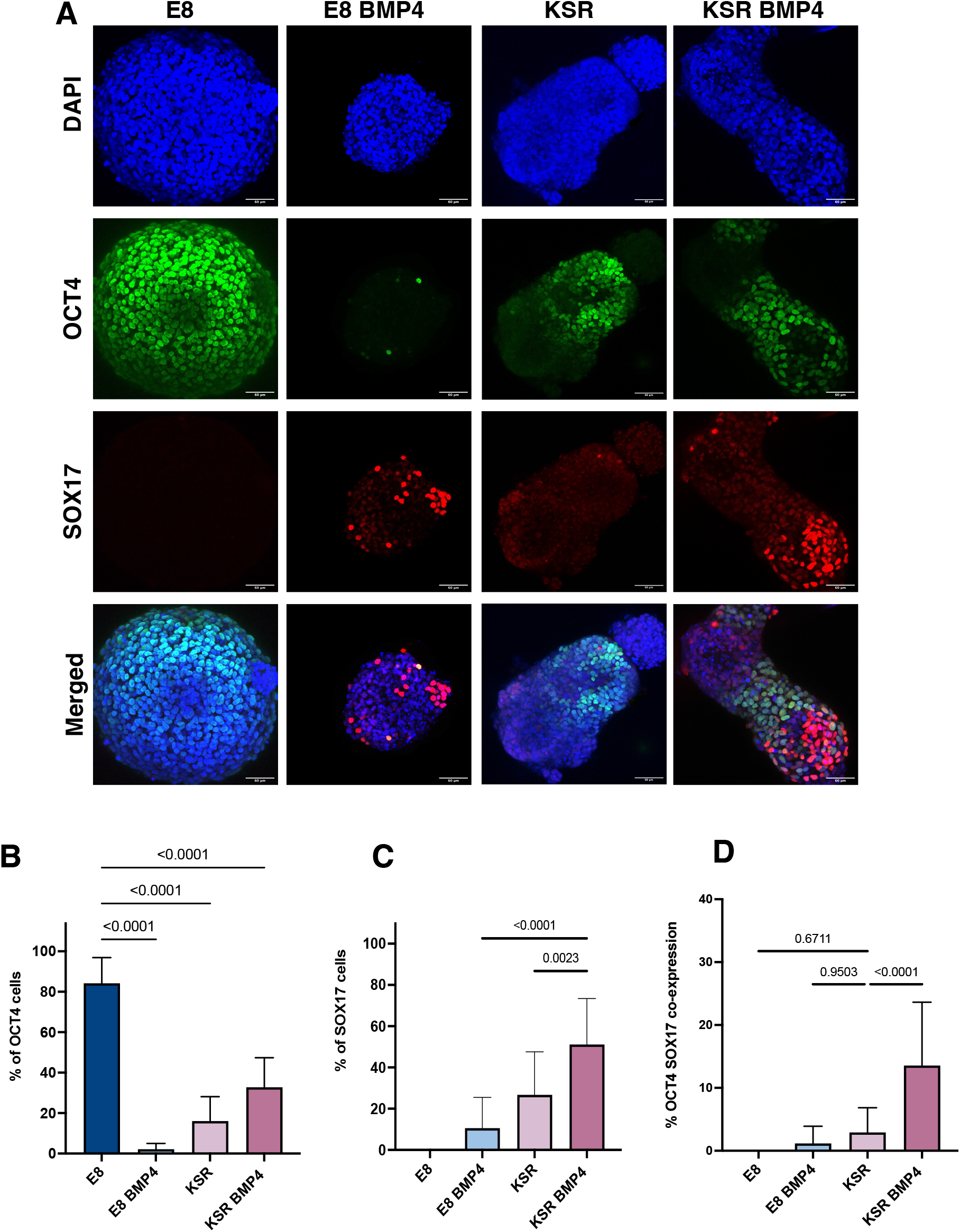
BMP4 signalling induces SOX17 specification in a 3D *in vitro* model of gastrulation. **A)** Immunostaining of OCT4 and SOX17 expression in each medium condition. Spheroids cultured in E8 medium maintained pluripotency OCT4 marker is highly expressed, while significant OCT4 reduction was observed in KSR BMP4 and both controls E8 BMP4 and KSR. Scale bars 60 μm. **B)** Quantification of the percentage OCT4 positive cells in E8 exhibits high OCT4 expression, while the low percentage of OCT4 positive cells in KSR BMP4, and both controls (n=17 spheroids, P-value < 0.0001, bars= mean value and SD). **C)** Percentage of SOX17 positive cells, the marker is not expressed in the E8 medium, while in KSR BMP4 shows the highest percentage of SOX17 expression. In control conditions, E8 BMP4 and KSR show less SOX17 percentage (n=13 spheroids, P-value <0.0001, bars= mean and SD). **D)** Quantification of the percentage of OCT4 SOX17 coexpressed cells under divers, highlighting the significant expression in KSR BMP4 (n =16, P-value <0.0001, bars= mean and SD).

We next interrogated markers for trilineage differentiation of germ layers. We stained for mesodermal markers BRA and EOMES and ectodermal marker SOX2 (see Supplementary S1-S2) to verify that 3D hiPSCs underwent a gastrulation-like process *in vitro*, which all showed the emergence of expression of lineage differentiation markers under diverse culture conditions. Nonetheless, we focussed on SOX17 as an apt readout and a proxy for lineage differentiation as it is expressed during the early stages of development and in the PE layer of the late blastocyst [9, 10]. SOX17 was absent in spheroids grown in E8; however, strong expression of SOX17 in KSR BMP4 conditions was detected consistently in the distal domain of elongated spheroids in a mutually exclusive expression pattern with OCT4 (Figure 3A). We also observed another cell population co-expressing SOX17^+^ /OCT4^+^ neighbouring SOX17^+ /^ OCT4^-^ cells in the elongated tip. We thus hypothesised that elongation of the spheroid and SOX17 expression could be in some way coupled. Consistent with this, when spheroids were cultured in E8 BMP4, low SOX17 expression was observed randomly. Some spheroids in this medium condition formed small protrusions, and SOX17 was expressed at a low level with marker polarisation in the protrusion area (Figure 3A). When spheroids were cultured in KSR medium, we observed lower expression of SOX17 (Figure 3C) that distributed homogeneously across the entire spheroid compared to the KSR BMP4 condition (Figure 3A). Altogether, these observations indicate a concomitant timing and suggest that morphogenesis and SOX17 marker polarisation toward the elongated tip in KSR BMP4 could be coupled.

Next, we quantified SOX17 expression in 3D using Imaris software to estimate the percentage of cells expressing markers (Figure 3C). Unsurprisingly, hiPSCs spheroids cultured in E8 medium maintained their capacity for self-renewal with high percentages of cells expressing OCT4 (Figure 3B). Conversely, a significant reduction of OCT4 expression was detected in all other medium conditions (E8 BMP4, KSR, and KSR BMP4) indicating cells ceased to be pluripotent. Spheroids in KSR BMP4 medium had high levels of SOX17 positive cells compared to other conditions (Figure 3C). We examined the expression of SOX2, an ectoderm marker, and mesodermal markers BRA and EOMES (Figure S1A-B; S2 A-C). Quantification showed that spheroids cultured in E8 medium express low levels of the ectodermal marker SOX2 with no sign of BRA expression. In the differentiating condition (KSR BMP4), SOX2 and BRA expression were observed (Figure S1A-B, S2A-B). When spheroids were grown in E8 BMP4 medium, neither SOX2 nor BRA was detected. Altogether, these observations demonstrate that in KSR BMP4 medium, spheroids give rise to cells that express markers for all three germ layers, whereas in KSR alone, spheroids undergo spontaneous differentiation mostly toward ectodermal lineage.

### PEG-peptide hydrogel encapsulation disturbs SOX17 patterning

When spheroids were exposed to BMP4, we observed the emergence of SOX17 expressing cells, which coincided with axial elongation. Therefore, we hypothesised that SOX17 expression was dependent on this shape change, and that impeding elongation would impact the expression of SOX17. To test this hypothesis, we physically confined spheroids using PEG-based hydrogels. We harnessed our fully synthetic platform that we previously reported could support the culture of hiPSC-derived intestinal organoids [28, 35]. These PEG-based hydrogels are formed at low polymer concentrations using two sequential reactions. First four-arm PEG-nitrophenyl carbonate (PEG-4NPC) is conjugated with a hetero-bifunctional peptide to form PEG-peptide conjugates. These conjugates are then crosslinked with four-arm PEG vinyl sulfone (PEG-4VS) to form the non-degradable hydrogel network (non-degPEG) (Figure 4A). We observed that confinement within non-degPEG prevented morphological changes in spheroids under differentiating conditions after BMP4 treatment (Figure 4B). Confined spheroids in E8 medium exhibited high levels of pluripotency marker OCT4, and no SOX17 expression was detected, similar to suspension conditions (Figure 4B, 6B). Interestingly, encapsulated spheroids in KSR BMP4 medium exhibited a significant reduction in OCT4 marker and low SOX17 expression (Figure 4B, 6C). In control conditions and in E8 BMP4 medium, we identified significantly reduced expression of OCT4 in addition to low expression of SOX17 (Figure 4B, 6C). In KSR medium, spheroids maintained high expression of OCT4, but no expression of SOX17 was detectable, which is comparable to the undifferentiated condition E8 (Figure 4B, 6C). Altogether, these results indicate that when cultured within confining hydrogels, cells within hiPSC spheroids do not express SOX17, as they do in free-floating differentiation conditions (Figure 3B-C). Overall, this suggests that embedding hiPSCs in a confining environment blocks morphogenesis, and despite the addition of BMP4, this was not sufficient to enable SOX17 expression.

**Figure 4.**
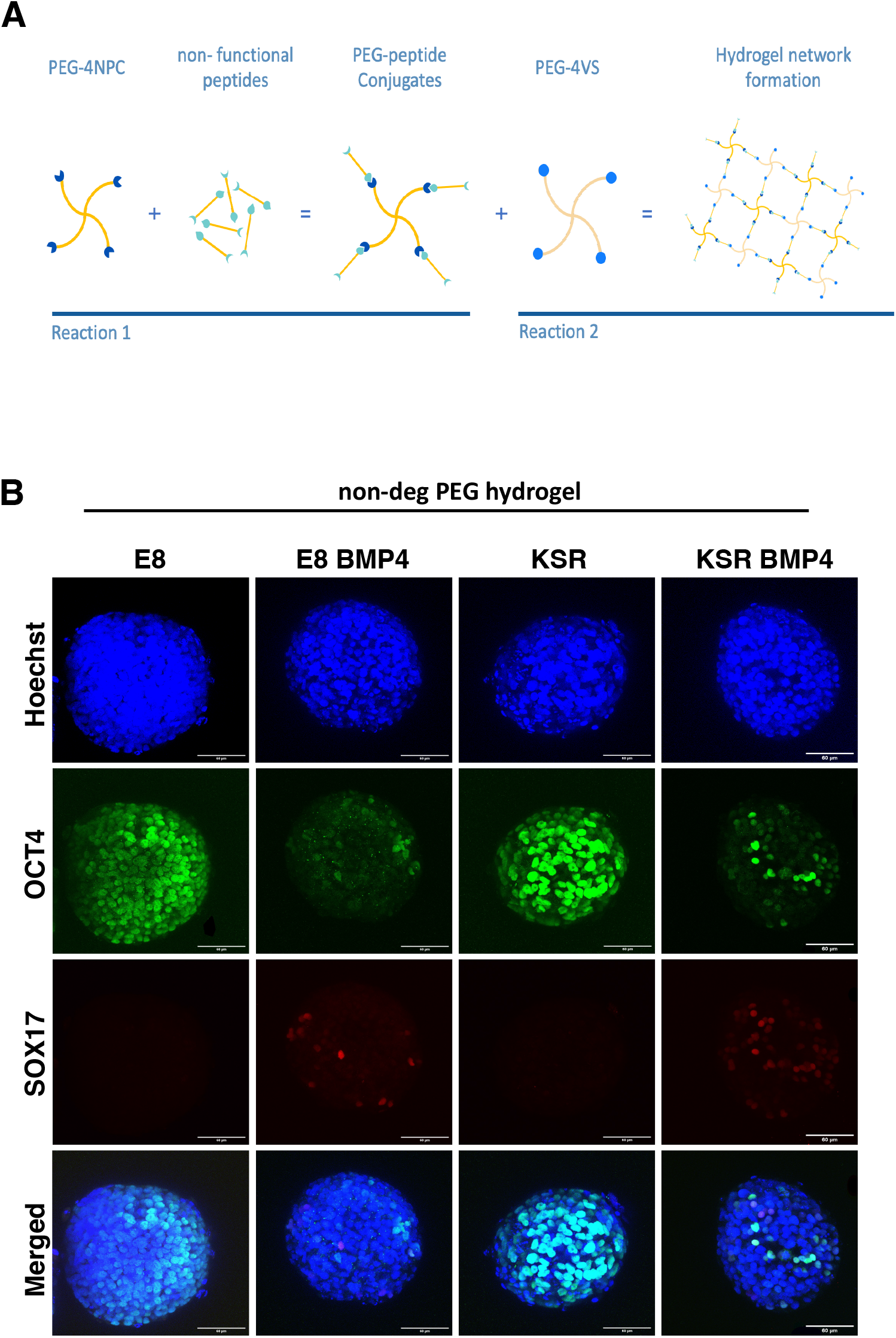
hiPSCs encapsulation in non-deg PEG hydrogel prevents morphological changes and decreases SOX17 expression. **A)** PEG-peptide hydrogel design based on two reactions, first PEG-4 NPC binds to a nonfunctional peptide (KDWERC) to form PEG-peptide conjugate with a solid content conc. 2.5%. This is followed by crosslinking with another PEG-4VS to form the hydrogel network. **B)** Immunostaining of OCT4 and SOX17. Spheroids encapsulated in non-deg PEG hydrogel cultured E8 medium maintained high OCT4 expression, whereas reduction in OCT4 was observed in KSR BMP4 and E8 BMP4 conditions, however, KSR exhibited high OCT4 expression. Scale bar 60 μm.

### Modulating PEG-peptide degradability promotes SOX17 expression

Induction of differentiation marker SOX17 was dramatically reduced compared to free-floating culture when spheroids were cultured in confining hydrogels. We thus asked whether a 3D hydrogel that was not confining would permit differentiation. We, therefore, modulated our design by swapping PEG-4VS with tetra-arm PEG molecules functionalised with acrylate groups (PEG-4Acr) to form a degradable hydrogel network (deg-PEG) (Figure 5A). PEG hydrogels cross-linked with acrylate have been reported to undergo controlled softening over time [26]. Using small amplitude oscillatory rheology, we confirmed that while standard PEG hydrogels cross-linked with PEG-4VS did not exhibit significant changes in stiffness over 5 days under standard culture conditions (G’ ~ 488.3 Pa to 329.9 Pa) (P value ns) (Figure 6A), hydrogels formed by swapping 75% of the PEG-4VS with PEG-4Acr softened over 5 days and were significantly softer than day 0 controls (~ 270 Pa to 78.7 Pa) (P < 0.0001) (Figure 6A).

**Figure 5.**
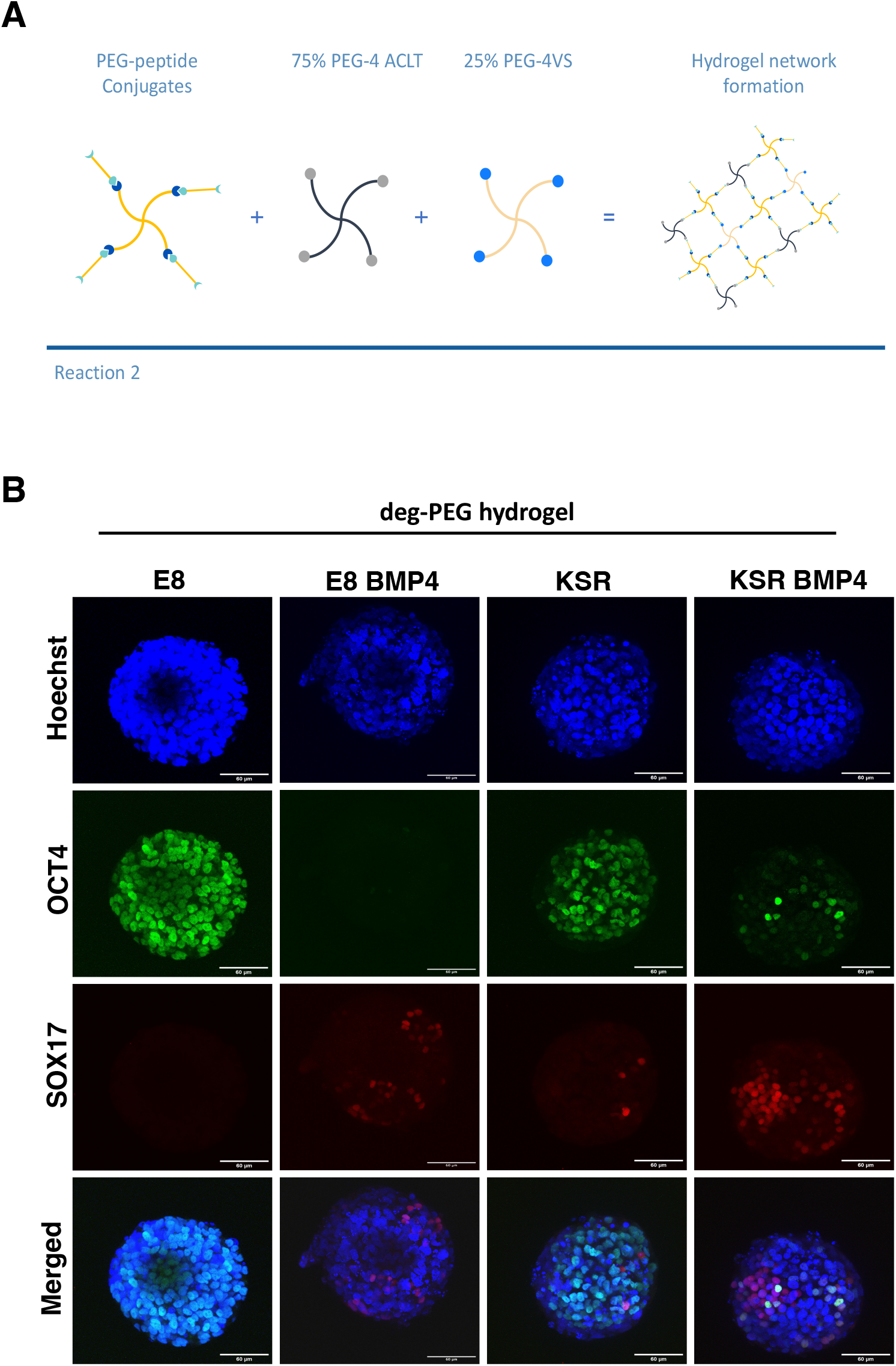
hiPSCs encapsulation in deg-PEG hydrogel prevents morphological changes and enhanced SOX17 expression. **A)** PEG-peptide modified hydrogel, here PEG-peptide conjugate is crosslinked with 25% PEG-4VS and 75% PEG-4ACLT to form the network. **B)** OCT4 and SOX17 markers, Spheroids embedded in deg-PEG hydrogel showed that OCT4 in E8 medium condition maintain the high expression of pluripotency, as expected in KSR BMP4 and E8 BMP4 the marker was reduced, and in KSR a decrease in the OCT4 expression was observed. Scale bar 60 *μ*m.

To determine whether encapsulation within deg-PEG hydrogels influences lineage differentiation, we stained for OCT4 and SOX17. Our findings show that in self-renewing conditions (E8 medium) encapsulated spheroids within deg-PEG maintained high expression of OCT4 and did not express SOX17 (Figure 5B). As expected, in KSR BMP4 medium condition we observed a significant reduction of OCT4 expression; we also identified an increase in SOX17 expression compared to non-degPEG (Figure 5B, 6B-C). Furthermore, when encapsulated spheroids were cultured in E8 BMP4, OCT4 remained downregulated albeit with a significant increase in SOX17 expression in contrast to E8 BMP4 in non-degPEG (Figure 5B, 6B-C). Spheroids grown in the KSR medium showed reduced expression of OCT4 in deg-PEG, when compared to non-degPEG conditions (Figure 6B) and SOX17 expression remained low (Figure 6C). Altogether these findings demonstrate that deg-PEG conditions partially restored SOX17 expression in both conditions KSR BMP4 and E8 BMP4.

### Biochemical cues regulate morphogenesis via cell proliferation and cellular tension

Next, we set out to explore whether biochemical cues, including BMP4 signalling, influenced the proliferation of free-floating hiPSC spheroids. This was investigated using an EdU proliferation assay on 3D spheroids in suspension. In E8 medium, cells were highly proliferative and exhibited homogenous EdU expression (Figure 7A-B). When spheroids were cultured in KSR BMP4 medium, cell proliferation was reduced and restricted to the tips of elongated spheroids (Figure 7A-B). In the intermediate conditions, the proliferation rate in E8 BMP4 was significantly decreased compared to E8, whereas spheroids cultured in KSR notably polarised toward the budding area (Figure 7A-B). Altogether, these results indicate that biochemical cues in the medium regulate cell proliferation, suggesting that low cell proliferation is associated with BMP4 addition, while KSR medium governs proliferative cell polarisation, which induces morphogenesis and differentiation.

**Figure 6.**
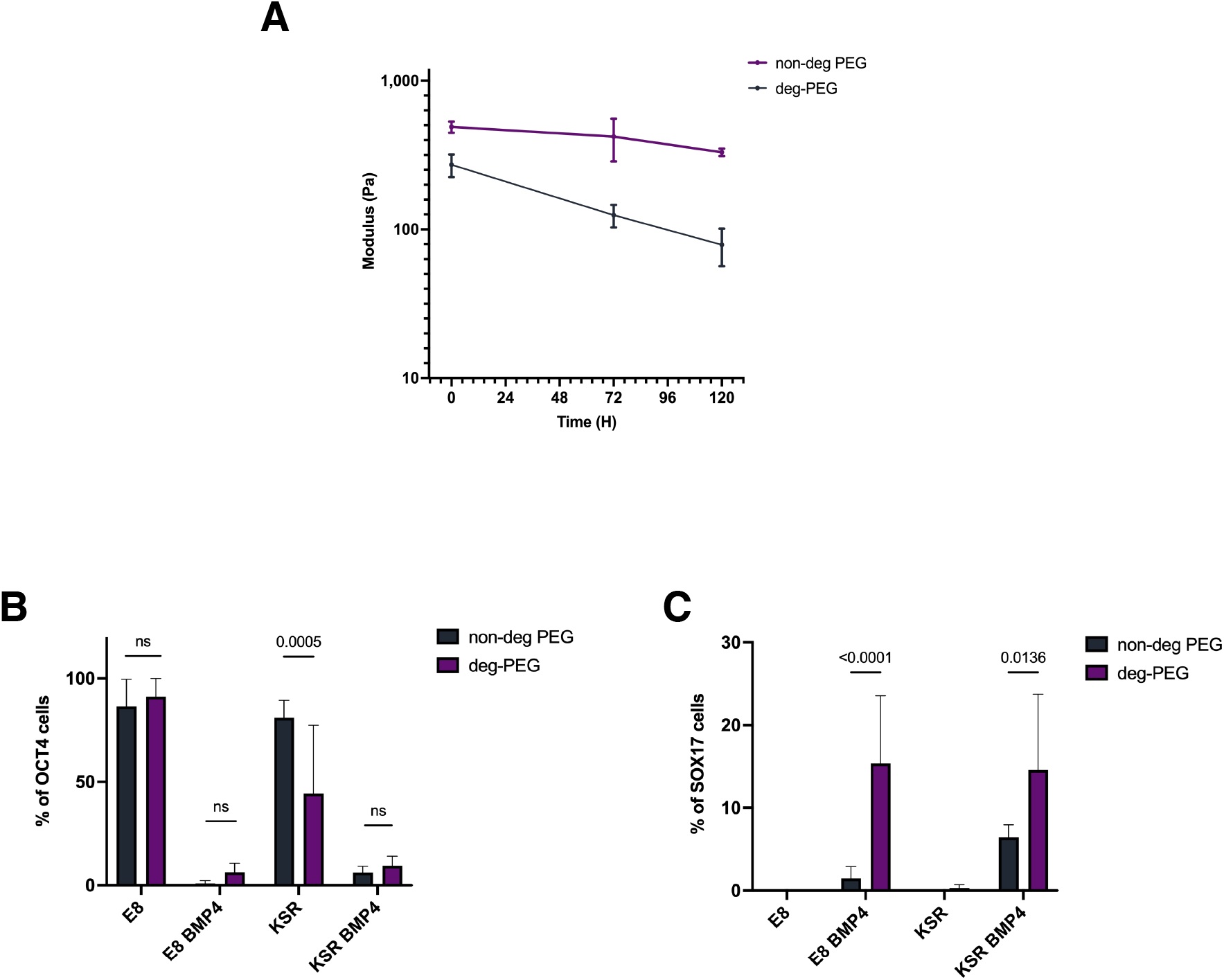
Diverse expression of OCT4 and SOX17 markers in non-deg PEG and deg PEG hydrogels. **A)** Rheology measurement shows the changes in hydrogel stiffness over time, the graph showed the high degradability in 75% PEG-4Acr (n=3 gels, bars SD). **D)** Quantification of tryptophan release in the media (n=3 gels, bars SD). **B)** Quantification of the percentage of positive cells expressing OCT4 in non-deg PEG hydrogel compared to deg-PEG hydrogel cultured in E8, KSR BMP4, E8 BMP4 and KSR (n= 8 spheroids per condition, P-value =0.0005, bars= mean and SD). **C)** Quantification of the percentage of SOX17-positive cells non-deg PEG hydrogel showing the significant reduction in KSR BMP4 and E8 BMP4 compared to deg-PEG hydrogel (n= 8 spheroids per condition, P-value <0.0001, P= 0.0136, bars= mean and SD).

**Figure 7.**
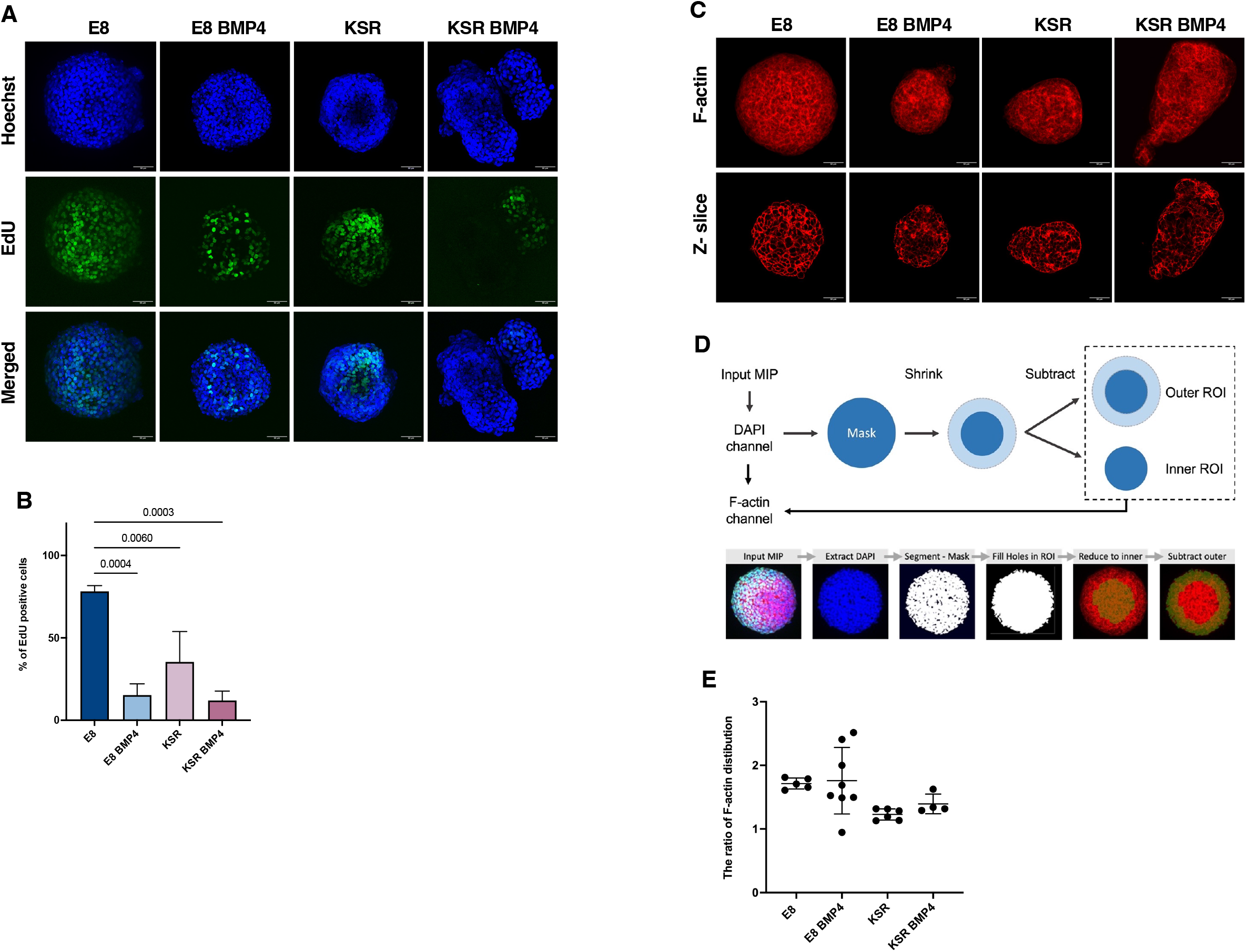
Distinct proliferation phenotype and F-actin distribution in response to biochemical cues. **A)** proliferation assay indicated by EdU staining shows high EdU positive cells in E8 medium, spheroids in KSR BMP4 show EdU positive cells restricted to the budding area. Both controls E8 BMP4 and KSR shows reduced EdU expression, in KSR medium EdU positive cells begin to polarise toward the budding. Scale bars 60 μm. **B)** quantification of the percentage of EdU expression, significant proliferation increases in E8 medium condition, in KSR BMP4 medium cells show lower proliferation rate. Controls E8 BMP4 and KSR exhibited a reduction in EdU expression (n=4 spheroids, P-value <0.0001, bars= mean and SD). **C)** Phalloidin staining shows a homogenous F-actin network in E8 medium condition, loose F-actin network was observed in KSR BMP4 condition, E8 BMP4 and KSR (z-slice; scale bar 60 μm). **D)** Schematic of a method to extract the average intensity of F-actin in the periphery and centre of gastruloids (top). Example of output images of a gastruloid after crucial steps of the analysis pipeline, displaying mask creation and ROI extraction (bottom). **E)** Quantification of the ratio of F-actin expression in the spheroid area, the graph shows the ratio of the distribution in the inner region, E8 condition exhibited consistent inner expression whereas, in KSR BMP4 less expression *is* shown in the inner area, in E8 BMP4 F-actin distribution varies and KSR condition displayed a decrease in the inner expression (n=4 spheroids).

Having observed variation in the number of proliferative cells post BMP4 treatment and under low FGF-b conditions (KSR), we next explored the F-actin network, which has been reported as an indicator of cellular tension [36]. Here, we postulate that cellular tension regulates spheroids’ morphology and can be inferred based on F-actin intensity and its localisation centrally or peripherally within spheroids (Figure 7 C-E). Phalloidin staining revealed that spheroids cultured in E8 medium exhibited organised and tightly packed F-actin with a homogenous network orientation (Figure 7C). In KSR BMP4 conditions, we observed a disorganised and stretched F-actin network localised preferentially around the edges (Figure 7C). However, in spheroids cultured in E8 BMP4, F-actin accumulated in the core of the spheroids (Figure 7C). In KSR medium, we also observed a disorganised F-actin network stretching around the edges of spheroids (Figure 7C). Overall, these observations show that changes in morphology correlate with changes in cellular tension (F-actin organisation) under distinct medium conditions and may suggest a causal relationship between the two.

Next, to validate our observation that morphological changes associated with cellular tension via F-actin expression, we applied an image analysis platform to quantify F-actin orientation; which calculates F-actin distribution based on the F-actin intensity and variations in its localisation (Figure 7D). F-actin orientation in round spheroids in E8 medium is mostly distributed in the inner core (centrally), however, in KSR BMP4, elongated spheroids showed more F-actin expression in the periphery (Figure 7E). In control conditions and E8 BMP4, F-actin was distributed in the centre, similar to E8 conditions, but with more variation between spheroids. Whereas in KSR conditions, F-actin was distributed in the periphery without significant differences compared to KSR medium conditions (Figure 7E). Altogether, these data indicate a relationship between the F-actin distribution and SOX17 expression.

## Discussion

The connection between form and function is a long sought-after problem in biology and beyond [37]. In early development, specification of three germ layers is controlled by morphogen signalling and mediated by physical forces between cells and the local tissue environment [2]. The physical properties of the native tissue vary (0.2-1 kPa) during developmental stages *in vivo* which regulate the self-renewing and differentiation process during early development [2, 38–41]. Nevertheless, the interconnection of patterning, cell fate and morphogenesis during human embryo development remains difficult to untangle. Human ‘gastruloids’ derived from hPSCs arrange as a multicellular system undergoing elongation and cells differentiate to form the three germ layers resembling the early stages of development [1, 42].

Here, we compare self-renewing versus differentiation conditions in suspension cultures of gastruloids, which give rise to morphological changes and the emergence of lineage specification. We then exploited PEG-based hydrogels by tuning their physical properties to interrogate whether physical confinement affects morphogenesis and differentiation [26]. In particular, we sought to investigate whether morphogenesis is required to obtain emergence of lineages in an organised manner. To address this question, 3D hiPSCs spheroids were embedded in PEG hydrogels with varying degradability. Our findings reveal that physical confinement prevented elongation, disturbing SOX17 expression. Modulating hydrogel degradability such that hydrogels soften over time re-established SOX17 expression, albeit impairing SOX17 polarisation when compared to 3D suspension conditions.

Our results demonstrate that biochemical cues trigger morphological changes. We robustly observed spherical shapes in E8, and axial elongation in KSR BMP4. In E8 BMP4, however, spheroids maintained a spherical shape while forming small buddings, and when spheroids were cultured in KSR, intermediate shapes formed undergoing spontaneous budding. High-content analysis approaches designed to obtain automated quantitative readouts from digital microscopy images provide multi-parametric data to quantify cell behaviours [43–45]. In this context, a robust high-content live imaging method was devised to monitor the 3D spheroids’ behaviour by quantifying changes in shape and area. Quantification and segmentation of the 3D spheroids’ phase contrast images are challenging and not often compatible with further analysis. However, we previously described a method using CellProfiler that can be easily applied in laboratories studying other biological systems to obtain an efficient analysis of 3D spheroids by quantifying area, size, shape, and roundness [34].

Several studies have investigated the role of the BMP4 signalling pathway in 2D micropattern systems [8, 16, 18]. However,in 3D models, ESCs treated with BMP4 were not able to elongate axially and differentiate toward the three germ layers [1, 23]. We have shown that 3D hiPSCs treated with BMP4 under defined culture conditions robustly break symmetry, elongate and generate tri-germ layers. Spheroids cultured in E8 medium remain pluripotent and express high level of OCT4. In contrast, cells in the KSR BMP4 spontaneously elongate mirroring A-P elongation and consistently present polarisation of SOX17 expression (endoderm) at the tip of the extended elongation.This suggests that in our model, BMP4 in KSR medium is sufficient to induce morphogenesis and germ layer specification, which is consistent with 2D micropattern models. Moreover, elongated spheroids exhibit OCT4 expression in the neighbouring elongated area of the spheroid. This is similar to the 2D micropattern in response to BMP4 treatment where hESCs expressed pluripotency markers in the colony centre. This phenotype resembles an early developmental phase when the primitive streak starts to form and the rest of the epiblast remains pluripotent [46].

Following observations suggesting that germ layer differentiation was associated with shape changes in free-floating culture, we aimed to impede this process using PEG hydrogels. We found that physical confinement prevents morphological changes and dramatically reduces the expression of SOX17. We then sought to identify if modulating hydrogel degradability could promote morphogenesis and differentiation. Deg-PEG hydrogels did not promote elongation, however, they did stimulate SOX17 expression in KSR BMP4 and E8 BMP4 conditions. Overall, this approach enables us to dissect the inter-correlation of 3D hiPSCs spheroids changes in shape with the patterning of germ layers.

During development, morphogenesis and cell fate decisions are regulated by the interplay between physical and morphogen signalling of the local environment [3, 47]. Cells generate tension via the cytoskeleton that drives changes in tissue morphology [47]. Here, we postulated that cellular tension impact spheroid morphology, and investigated F-actin distribution as a readout for these forces. F-actin in the E8 medium condition was distributed centrally suggesting that tension is homogeneous and perhaps drives the sphere to expand. In the KSR BMP4 condition, F-actin was distributed around the periphery, indicating higher tension around the edges, especially in the elongated area. In the E8 BMP4 condition, F-actin staining showed a central distribution and accumulation of F-actin adjacent to buds. These observations suggest that in both differentiation conditions (KSR and KSR BMP4), cellular tension might contribute to morphogenesis and is in keeping with reports that the polymerisation of the F-actin network toward the plasma membrane generates pushing forces resulting in shape changes [36].

This work focussed on investigating how biochemical cues and physical confinement separately influence morphogenesis and germ layers specification in 3D hiPSCs-derived models of gastrulation. These outcomes open opportunities for further investigation to characterise cell lineages expressed upon BMP4 treatment, and to explore physical forces generated by cell-cell adhesion or actomyosin contractility. Finally, our work also offers building blocks towards hiPSCs-based high-throughput platforms to screen for cellular responses to chemicals, genomic perturbations or genetic backgrounds to early development.

## Methods

### hiPSCs culture

hiPSCs line (Hoik_1) were selected from HipSci biobank (www.hipsci.org). As previously described [48] hiPSC were derived from skin fibroblast and reprogrammed using Sendai virus vectors (CytoTune) expressing the four factors OCT4, SOX2, MYC and KLF4. All cell lines have been assessed for quality control profile to verify pluripotency and genotyping assay. Briefly, iPSCs were cultured on a previously coated 6-well plate (Greiner) with 10 ug/ml vitronectin (Stem cell Technologies). Cells were cultivated in feeder-free medium using Essential 8 (E8) medium with 2% E8 supplement (50x) (Thermo Fisher Scientific), and 1% Penicillin Streptomycin (5000U/ml) (Gibco) incubated at 37 ° C, 5% CO_2_. Routinely, hiPSCs required daily feeding and passaging every 4-5 days when it reached 80% confluence.

### Spheroids derivation from hiPSCs

To dissociate hiPSCs colonies into single cells, colonies were washed with Hank’s Balanced Salt solution HBSS (Gibco). Cells were incubated in Accutase (Bio Legend) at 37° C, 5% CO_2_ for 4 minutes and resuspended in E8 medium (Thermo Fisher Scientific) and 10 μM Y-27632 Rho-kinase inhibitor (ROCKi) (ENZO Life Sciences). Prior to cell seeding, low attachment 96-well V bottom plates (Thermo Fisher Scientific) were coated with 5% (w/v) Pluronic solution (Sigma). Plates were centrifuged at 500 x g for 5 min and incubated at room temperature (RT) for 30 min, followed by washing with Phosphate Buffer Saline PBS (Gibco). Single cells were seeded at a density of 750 cells/ well and cultured in E8 medium (Thermo Fisher Scientific) as described previously and 10 μM ROCKi (ENZO Life Sciences) to prevent cell apoptosis and enhance cell viability. hiPSCs were incubated for a further 48 hrs at 37° C, 5% CO_2_ to form spheroids.

### Gastrulation-like induction of 3D hiPSCs

hiPSCs spheroids were cultured in self-renewing and differentiation conditions at 37° C, 5% CO_2_ for 96 hours, as follows; (1) E8 medium supplemented with 2% E8 supplement (50x) (Thermo Fisher Scientific), 1% Penicillin Streptomycin (5000U/ml) (Gibco), and 10 μM ROCKi (ENZO Life Sciences); (2) Knock out serum medium KSR consisting of Advance DMEM/F-12 medium, supplemented with 20% KnockOut serum replacement, 1% L-Glutamine, 1% Penicillin Streptomycin (5000U/ml) (all Gibco), 0.1mM β-mercaptoethanol (Sigma), 10ng/ml basic fibroblast growth factor (bFGF), 50ng/ml BMP4 (Invitrogen), and 10 μM ROCKi (ENZO Life Sciences). Control medium (3) E8 medium composition supplemented with 50ng/ml BMP4; and (4) KSR medium as described above without 50ng/ml BMP4..

### Live imaging

hiPSCs spheroids cultured in E8 medium, KSR BMP4 medium, E8 BMP4 medium and KSR medium were imaged for 96 hrs using JuLI™ Stage Real-Time Cell History Recorder (NanoEnTek). Time lapse images were acquired every hour.

### Hydrogel fabrication

Hydrogels were synthesised as previously described [28, 33] using 4-arm PEG-peptide conjugates and 4-arm PEG-VS 20 kDa (non-degradable) or a mixture of 25% 4-arm PEG-VS and 75% 4-arm PEG-ACLT 20 kDa (degradable). Briefly, peptide conjugates were synthesised with Ac-KDW-ERC-NH2 (custom synthesis Peptide Protein Research, Ltd. (UK), >98% purity) and 4-arm 10 kDa PEG-NPC (JenKem Technology, USA). The N-terminal primary amine (lysine) of Ac-KDW-ERC-NH_2_ was reacted with PEG-NPC to form conjugates. Purified conjugates were crosslinked with either PEG-4VS or PEG-4 ACLT to form the hydrogel network through a Michael addition. Here, the reaction between peptide conjugate and crosslinkers (PEG-4VS and PEG-4 ACLT) was performed in stoichiometric ratio 1:1 in30mM HEPES buffer at pH 8 (Sigma) diluted in 1x HBSS (Gibco). Hydrogels were allowed to form for 30-45 min t. The solid content of the polymer was 2.5% (w/v) for both hydrogel conditions.

### Rheological measurements of hydrogels

Hydrogel storage modulus G’ was measured on a strain-controlled ARES from TA Instruments using an 8 mm plate with a 0.01-rad angle. 50 μL samples were prepared in 8mm glass rings, placed onto the rheometer plate, and measurements carried out at 37 °C while sealed with paraffin oil to prevent evaporation. A frequency sweep was recorded, measuring *G’* as a function of shear frequency in the range 100–0.1 rad s–1 at a fixed strain of 1%. (Orchestrator software, version 7.2.0.2).

### hiPSCs spheroids encapsulation in PEG-peptide hydrogel

hiPSCs spheroids were obtained 48 hrs post seeding in 96 wells V-bottom plate as previously described. The spheroid in each well was washed once with 30mM HEPES buffer (pH 8.0), then resuspended in 30mM HEPES buffer (pH 8.0) (Sigma). Spheroids were encapsulated within 10 μl of hydrogel, formed as described above. in μ-slides angiogenesis glass bottom (ibidi) by placing the mixture directly in each well and incubating for 45 min at 37° C, 5% CO_2_. Medium containing 10 μM ROCKi (ENZO Life Sciences) was then added and cells were cultured for 96 h a at 37° C, 5% CO_2_ with daily feeding.

### Immunofluorescence staining

Spheroids in suspension were fixed using 4% paraformaldehyde (PFA) (Sigma) incubated at room temperature (RT) for 45 min on a shaker followed by 3x washing with PBS (Gibco). Cells were permeabilised and blocked with 0.3% Triton X100 (Sigma) in PBS (Gibco), and 3% bovine serum albumin (BSA) (Sigma) for 1 h at RT on a shaker. Spheroids were stained with primary antibodies: rabbit anti-OCT4 (Abcam, 1:500); goat anti-SOX17 (R&D, 1:100), goat anti-SOX2 (R&D, 1:200); goat anti-Brachyury (R&D, 1:100); and rabbit anti-Eomes (Abcam,1:100) were diluted in the blocking buffer and incubated it overnight in dark at 4°C on a shaker. Cells were washed 3X with 0.1% TritonX100 (Sigma) in PBS (Gibco) followed by secondary antibodies all (Thermo Fisher Scientific, 1:500) Alexa fluor 488 anti-rabbit, Alexa fluor 633 anti-goat, Alexa fluor 594 anti-mouse, and Alexa fluor 555 Phalloidin (Thermo Fisher Scientific, 1:100), and DAPI (Invitrogen, 1:1000) were added and incubated for 2 h at RT on shaker followed by 3x washes in 0.1% TritonX100 (Sigma) in PBS (Gibco).

Encapsulated spheroids in PEG hydrogels were fixed with 4% PFA (Sigma) incubated at RT for 45 min on a shaker and washed 3x with PBS (Gibco) 10 min each. Cells were permeabilised and blocked with 0.3% Triton X100 (Sigma) in PBS (Gibco) for 1 h followed by a blocking buffer consisting of 0.1% TritonX100 (Sigma) plus 3% BSA (Sigma) and were incubated overnight at 4°C on a shaker. Primary antibodies: rabbit anti-OCT4 (Abcam, 1:50); goat anti-SOX17 (R&D, 1:50) were diluted and YAP 1 (Santa Cruz, 1:50) were diluted in the blocking buffer and incubated for 48 h in the dark at 4°C on a shaker. This was followed by 5x washing using 0.1% TritonX100 (Sigma) in PBS (Gibco) for 10 min each. Spheroids were stained with secondary antibodies all (Thermo fisher, 1:50) Alexa fluor 488 anti-rabbit, Alexa fluor 633 anti-goat, Alexa fluor 594 anti-mouse, and Alexa fluor 555 Phalloidin (ThermoFisher, 1:50), and DAPI (Invitrogen, 1:1000) overnight in the dark at 4°C with shaking. Cells were then washed 5x with 0.1% TritonX100 (Sigma) in PBS (Gibco) for 10 min each.

### Cell proliferation assay

Spheroids in suspension at day 4 were labelled with 4uM EdU and incubated for 6 h using Click-iT EdU Alexa fluor 488 imaging kit (Thermo Fisher Scientific) following the manufacturer’s protocol.

### Imaging and statistics

Images were acquired using a Leica TCS SP8 confocal microscope with 40x oil objective to image spheroids in suspension. Encapsulated spheroids in μ-slides angiogenesis glass bottom (ibidi) were imaged with 63x oil objective. Image analysis was obtained using Imaris software version 9.9, and statistical analysis was performed in GraphPad Prism version 9.3.1. Ordinary one-way ANOVA test for multiple comparisons (Sidak’s) followed by a post-hoc Tukey test.

## Supplemental information

Supplementary figures S1, and S2

Movies S1,S2,S3, and S4.

## Acknowledgements

We thank the Ministry of Education in Saudi Arabia for the PhD studentship and for funding this project by the Saudi Arabian Cultural Bureau in the UK. EG acknowledges funding from the EPSRC (EP/V04723X/1). We gratefully acknowledge Prof Fiona Watt for her support and valuable discussion. Special thanks to Dr Mukul Tewary, Dr Alice Vickers, James Williams, Lazarous Fotopoulos, and Fuad Mosis.

## Author contributions

D.D, E.G, supervision, writing review and editing. H.A, D.D, conceptualization, funding acquisition, investigation, data curation, writing-original draft, methodology, project administration. H.A., E.R. formal analysis. T.W, H.A visualisation, imaging acquisition. A.K, Y.G hydrogel experiments. J.G. writing-review and editing.

---

**Figure S1. The emergence of mesodermal markers BRA and Eomes.in 3D *in vitro* model of gastrulation.**

**A)** Immunostaining of BRA and Eomes expression. In E8 medium maintained mesoderm differentiation was not detected, in KSR BMP4 induced the expression of BRA with a slight expression of Eomes marker, E8 BMP4 showed very low Eomes expression, whereas in KSR cells exhibited low BRA expression. Scale bars 60 μm. **B)** Quantification of the percentage BRA positive cells in KSR BMP4, and KSR (n=13 spheroids, P-value = 0.0175, bars= mean value and SD). **C)** Quantified percentage of Eomes positive cells in KSR BMP4 showed reduced expression, similarly in E8 BMP4 and KSR (n=13 spheroids, P-value = ns, bars= mean and SD).

**Figure S2. The specification of ectodermal marker SOX2 in 3D *in vitro* model of gastrulation**

**A)** Immunostaining images representing the OCT4 (pluripotency) and SOX2 marker expressions. OCT4 marker is highly expressed in the self-renewing medium E8, a significant decrease in OCT4 expression was observed in KSR BMP4, E8 BMP4 and KSR. Scale bars 60 μm. **B)** Quantification of the percentage SOX2 positive cells in E8 exhibits low SOX2 expression, in KSR BMP4 moderate SOX2 expression, while the marker was not observed in E8 BMP4, and both controls (n=8 spheroids, P-value < 0.0089, bars= mean value and SD). **C)** The percentage of positive cells co-expressing OCT4/SOX2 marker, which is shown in KSR BMP4 and KSR (n=7 spheroids, P-value <0.0001, bars= mean and SD).

